# Pathway analysis of GWAS loci identifies novel drug targets and repurposing opportunities

**DOI:** 10.1101/425769

**Authors:** Deepali Jhamb, Michal Magid-Slav, Mark R. Hurle, Pankaj Agarwal

## Abstract

Genome-wide association studies (GWAS) have made considerable progress and there is emerging evidence that genetics-based targets can lead to 28% more launched drugs. However, translating the results of GWAS for drug discovery remains challenging. We analyzed 1,589 GWAS across 1,456 protein interaction pathways to translate these often, imprecise genetic loci into therapeutic hypotheses for 182 diseases. We validate these pathway-based genetic targets by testing if current drug targets are enriched in the pathway space of the same indication. Remarkably, 30% of diseases have significantly more targets in these pathways than expected by chance; the comparable number for GWAS alone (without using pathway analysis) is zero. Although pathway analysis is routine for GWAS, this study shows that the routine analysis can often enrich for drug targets, by performing a systematic global analysis to translate genetic findings into therapeutic hypotheses for new drug discovery and repositioning opportunities for current drugs.

## Introduction

Decades of research into the genetic basis of human disease has shown that most common diseases are highly polygenic in nature and hence understanding the pathways or the mechanisms by which these genes interact at a pathway or systems level and exert biological effect is essential. Previous studies have shown that GWAS genes are enriched for known drug targets (Nelson et al, 2015; Sanseau et al, 2012); however, most of the GWAS targets are not druggable using either small molecules or biopharmaceuticals. In contrast to a single GWAS locus, single gene perspective, we hypothesized that integrating GWAS loci with biological pathways could add value for drug discovery by identifying drug targets and novel pathways enriched in genetic risk loci.

GWAS has made substantial progress in identifying susceptibility loci associated with complex traits and diseases (Visscher et al, 2017). Pathways (Hirschhorn, 2009; Wang et al, 2010), networks (Cao & Moult, 2014; Sun et al, 2015), gene expression (Pers et al, 2015), literature mining (Ailem et al, 2016; Raychaudhuri et al, 2009), and gene set enrichment (Mooney et al, 2014) have been used to determine the functional relevance of GWAS variants. We hypothesized that the connection between GWAS hits and drug targets might be more easily discernible in pathway space as it is likely that both disease genetics and drug targets for a disease point to the same pathways. Published pathway analysis has concentrated mainly on individual or a small group of GWAS; the one large scale analysis was aimed at linking GWAS phenotypes based on shared KEGG pathways (Brodie et al, 2014).

## Results and Discussion

To begin, we repeated an earlier analysis (Sanseau et al, 2012) to see if all the significant GWAS genes (see Materials and Methods) in the pathway universe are overall enriched for drug targets. We observed 285 GWAS genes that are drug targets (expected=217, odds ratio = 1.31 (1.12-1.55), two-sided Fisher’s exact test, p=6.2e-9). Thus, GWAS genes within the pathways are indeed enriched for drug targets. However, a major limitation of this calculation is that it does not consider if the GWAS traits match the indication for these drug targets. If we limit the analysis to the GWAS trait matching the drug indication, the enrichment is not detectable (Supplementary Table 1). An earlier study has detected enrichment across disease classes if both Mendelian and common disease genes are grouped and related diseases are considered as matched (Nelson et al, 2015).

To investigate our hypothesis, we first obtained 1,589 GWAS from STOPGAP, a comprehensive resource of genetic associations (Shen et al, 2017). We processed the data using the pipeline summarized in Fig. 1. A crucial step in interpreting GWAS results is to map variants to their effector genes. Causal genes for a GWAS loci are frequently identified using expression trait loci (eQTLs) in disease relevant tissue or coding variants in high LD. Unfortunately, only a small percentage of causal genes can be identified with these methods because eQTL data is not available for many cell types and few loci have missense coding variants (Spain & Barrett, 2015). In addition, the causal genes picked by these methods are not always accurate. To avoid the bias of picking a single effector gene, we considered all “candidate effector genes” (Shen et al, 2017) associated with all variants in high linkage disequilibrium having an r^2^≥0.7. Overall, we obtained 13,172 genes associated with 278 traits. During pathway enrichment, for each trait we adjusted the number of genes mapped from a variant to ensure that at most one gene from each locus was counted (see Materials and Methods) (Agarwal et al, 2006). This lenient variant to gene mapping assumes that use of pathway information will implicitly enrich for the correct GWAS target.

**Figure 1:**
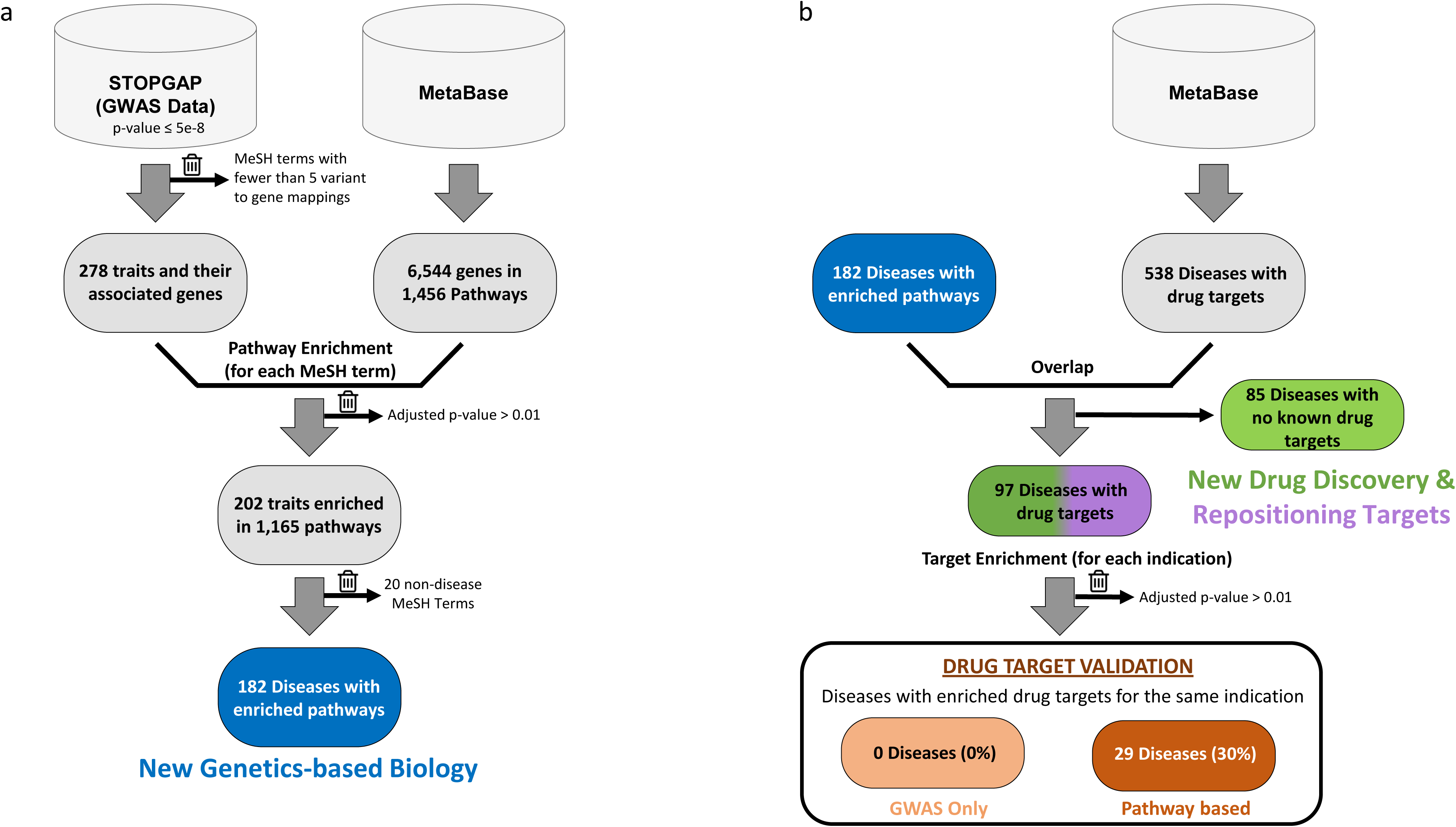
Overview of the workflow for data processing. Method used for (a) pathway enrichment and (b) drug target validation. The trash icon indicates items that were excluded from any further analysis.

For the pathway enrichment, MetaBase (Nikolsky et al, 2009) was used to obtain a collection of 1,456 manually curated pathways. These pathways include the major functional categories of cell signaling and human metabolism. In our analysis, we identified 202 of the 278 GWAS linked traits enriched for pathways at FDR p<0.01 (Supplementary Table 2). Of these 202 traits, 20 were not diseases and were excluded from further analysis. A majority of these 182 diseases show a significant enrichment for 10 or less pathways (Supplementary Fig. 1). Some of the diseases however, related to inflammation, digestive system, and metabolism are associated with many significant pathways, possibly highlighting the role of several mechanisms that contribute to the development of these diseases (Supplementary Fig. 2).

We further examined the 182 diseases by asking whether the genetically-identified pathways were enriched for drug targets for the same indication (Fig. 1b); for example, if drug targets in the pathways identified using asthma GWAS genes are indeed drug targets for asthma. Only 97 of these 182 diseases, however, have one or more known drug targets as defined in MetaBase (Nikolsky et al, 2009). We constructed a 2×2 contingency table for each of these 97 diseases, and tested for drug target enrichment in the pathway space. Remarkably, for 29 (30%) of these 97 diseases, the pathways were enriched in drug targets for the same indication at FDR p≤0.01 (Fig. 2b, Supplementary Table 3). In contrast, 0 (0%) of these 97 diseases show enrichment without the pathway step i.e. GWAS alone (Fig. 2a). Hence, mapping GWAS target genes into pathways reveals a concordance of GWAS targets within pathways containing already successful drug targets.

**Figure 2:**
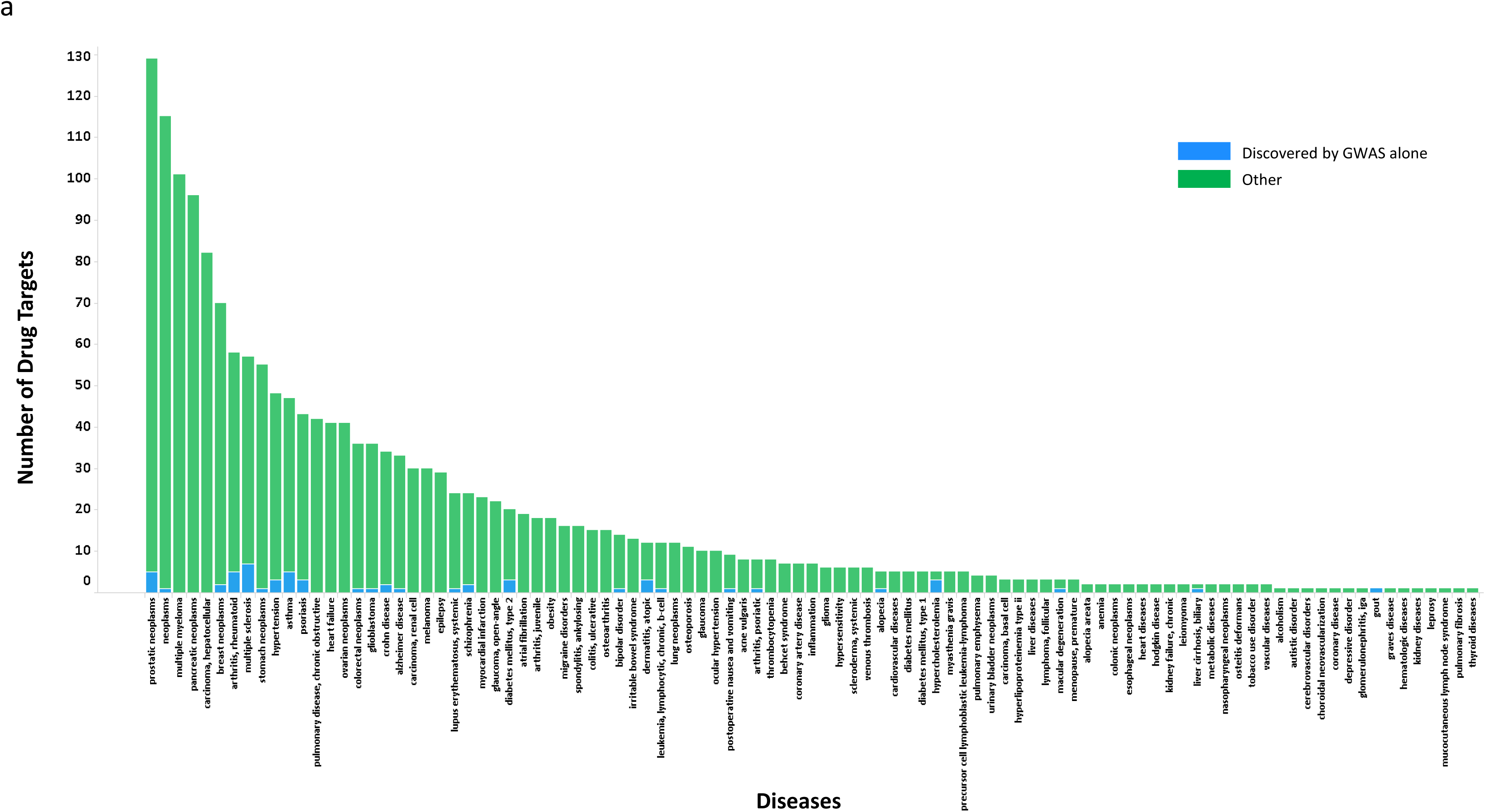
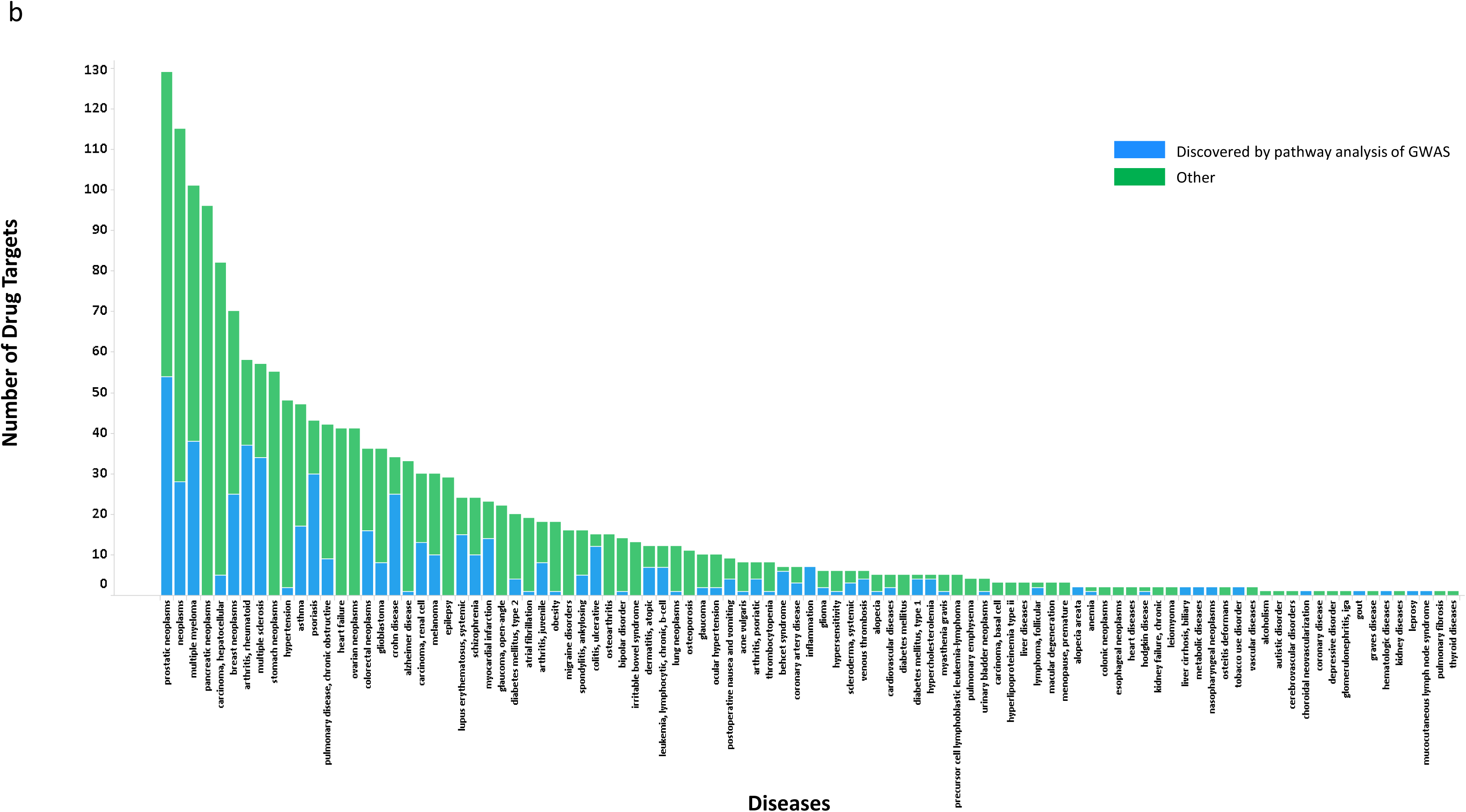
Pathway based Drug Target Enrichment for GWAS. (a) Number of drug targets discovered by GWAS data alone (b) Number of drug targets discovered by pathway analysis of GWAS. The data is shown for each of the 97 diseases with drug targets. The difference in blue between (a) and (b) represents the value of pathway analysis.

As the pathway approach adds value, we can pursue new genetically-identified biological targets in these 182 diseases, whether (for 97 of them) or not (for 85 of them) there is a drug target against the disease. This suggests that pathway analysis can identify multiple GWAS traits with actionable pathways from both repurposing and novel target discovery perspectives. As an example, we closely examined the results for a disease with a small number of drug targets. Behçet’s is a chronic autoimmune and auto-inflammatory disease that can affect almost all organ systems and is associated with increased mortality and high morbidity(Nair & Moots, 2017). For example, a survey of the top-ranked GWAS genes from the 19 variants associated with Behçet’s syndrome would find two drug targets, IL12A and CCR1, for diseases other than Behçet’s, but it would find none of the seven current Behçet’s drug targets. The pathway approach, however, finds six of these seven Behçet’s drug targets in the 12 pathways containing an overrepresentation of genes from the 19 loci (Supplementary Fig. 3).

The non-Behçet’s drug targets found in these 12 pathways represent the repositioning opportunities that are most aligned with the GWAS findings. As an example, IL12B is a member of many of these 12 pathways, but is not in a GWAS locus. Thus, we hypothesized that ustekinumab, a monoclonal antibody that blocks IL12 and IL23, could benefit Behçet’s patients. As it turns out, it is currently being clinically tested for Behçet’s s disease in an open-label trial (ClinicalTrials.gov), though this was not recorded in MetaBase. It is also noteworthy that six of the 19 Behçet’s GWAS loci are represented in these 12 pathways and these loci include IL12A, its receptor subunit IL12RB2 and the downstream signaling molecules STAT4 and IL10. Further, pathways which do not include one of the seven current Behçet’s targets may represent interesting novel biology.

This analysis can also identify correlated diseases with shared biological mechanisms. For example, “Altered Ca2+ handling in heart failure” pathway is significantly associated with GWAS genes from both tachycardia and autistic disorder, suggesting that the drugs associated with tachycardia could potentially be used for treating autism. A recent review article examined several reports on the use of propranolol, a beta-adrenergic antagonist, and indicated that this drug can help manage emotional, behavioral, and autonomic dysregulation in autism spectrum disorder (Sagar-Ouriaghli et al, 2018).

There are a few caveats with our analysis. First, the pathway collection includes canonical signaling pathways and other useful constructs in terms of disease biology, though these are likely biased in terms of emphasizing known biology, but hopefully it is not through excessive inclusion of potential GWAS genes. Second, GWAS, by itself, does not provide the required direction of modulation for the target (increase or decrease), i.e. whether we need an agonist or antagonist for therapeutic benefit. Thus, the identified direction for repurposing could be wrong. However, often the current indication of the drug will be helpful. For example, a current anti-inflammatory in Behçet’s pathway is likely to have the correct direction for Behçet’s as well. Third, we also took an optimistic view of each genetic locus preferring genes related through pathways as most likely suspects. We also performed the analysis by taking the “best” gene in each locus (independent of pathway) and observed a significant though reduced enrichment for pathways (see Materials and Methods).

Previously published studies have shown that progressing genetically validated targets can significantly increase the chances for clinical success (Cook et al, 2014; Nelson et al, 2015) and that genetics-based targets can lead to 28±18% more launched drugs (Hurle et al, 2016). This global pathway analysis of GWA studies provides a framework to translate common disease genetics into therapeutic hypotheses, and thus realize that 28% gain in productivity. Overall, our systematic analysis of GWAS enriched pathways identified known and novel pathways with drug targets for 182 diseases. This provides mechanistic pathway hypotheses for each of these diseases with multiple tractable targets in each pathway and drug repositioning opportunities for many diseases. We validated our findings by showing that 30% of the diseases have significantly more targets in these pathways than expected by chance, which is remarkable considering that none of these diseases were enriched for targets using GWAS alone. This highlights the potential for drug discovery to focus the search for novel targets and repositioning opportunities within biological pathways enriched for GWAS targets.

## Materials and Methods

### GWAS Data Processing

The GWAS data was obtained using STOPGAP v2.5.1 (Shen et al, 2017). STOPGAP includes data from NHGRI, GWASDB, and GRASP, which we filtered to retain only significant variants with p-value ≤5e-8, resulting in a set of 473 traits obtained from 1,589 GWA studies. We defined traits based on the MeSH classification available from STOPGAP and the same definition of traits is used throughout the paper. Further, any records containing genes that did not map to Entrez ids were removed and any traits with fewer than five variant to gene mappings were also eliminated. This yielded a final set of 278 traits derived from 1,398 GWA studies. In order to avoid the biases of picking a single methodology to assign an effector gene, we included all “candidate effector genes” (Shen et al, 2017) associated with all variants in high linkage disequilibrium having an r^2^≥0.7. Overall, we obtained 13,172 genes associated with 278 traits, but we subsequently corrected for the number of genes mapped to a variant during pathway enrichment to ensure that at most one gene from each locus was counted. As an example, if four genes on a pathway were obtained from the same GWAS locus for that disease, we only included one of those genes for enrichment calculations to avoid erroneous results. This approach takes an optimistic pathway view of each variant-to-gene mapping. All these “candidate effector genes” associated with these traits were extracted and used for pathway enrichment. To determine if the GWAS genes in pathways are overall enriched for drug targets (as in paragraph 2 of the main section), we carried out the pathway enrichment for 5,305 “best” GWAS genes (as described in STOPGAP) associated with 261 traits (data not shown). The data for “best” genes was processed in a similar fashion to all “candidate effector genes”.

### Pathway Enrichment

Pathway information was obtained from MetaBase version 6.30.68780. MetaBase, by Clarivate Analytics, is the manually curated knowledge base behind MetaCore, a software suite for pathway and network analysis of high throughput sequencing data (Nikolsky et al, 2009). It contains 1,456 protein interaction pathways, which are a comprehensive resource of human, mouse, and rat signaling, metabolism, diseases, and stem cells. R programming language was used to query the database and retrieve enriched pathways. The significance of enrichment was assessed by p-values of hypergeometric distribution (Nikolsky et al, 2009). Our null hypothesis was that there is no significant pathway enriched for a given trait. Benjamini-Hochberg method was used to obtain the adjusted p-values and a threshold of FDR corrected p-value ≤0.01 was used to determine the significance (Supplementary Table 2).

### Drug Data and Enrichment

#### Drug data

Drug information for already launched drugs or those which are in preclinical or clinical development was obtained from MetaBase version 6.30.68780. 672 proteins (gene identifiers) had active projects targeting them. These drugs are for 538 indications which were mapped to Diseases within the MetaBase resource.

#### Enrichment analysis

All enrichments were computed using a Fisher Exact test using 2×2 contingency tables with a Perl module (Lanczos, 1964). A total of 7,152 genes are present on at least one of the 1,456 MetaBase pathways and of these only 6,544 mapped to HUGO symbols for protein coding genes and we used this as our pathway universe to obtain enrichment of drug targets for a matching indication. This is the universe that was used for the calculation in paragraph 2 of Main section.

We computed a 2×2 contingency table to check if all the GWAS genes in pathways are overall enriched for drug targets.

|  | Drug Target | Not Drug target | Total |
| --- | --- | --- | --- |
| GWAS gene | 296 | 1,926 | 2,222 |
| NOT GWAS gene | 376 | 3,946 | 4,322 |
| Total | 672 | 5,872 | 6,544 |

The expected number of GWAS genes that are drug targets in these pathways is 216.88 the observed number of 285 with an odds ratio of 1.31 (1.12-1.55) with p=6.2e-9. Thus, GWAS genes within this pathway universe are enriched for drug targets.

We further checked if the enrichment (FDR p<0.01) holds up for each disease, and built contingency tables for each MeSH disease. For example, for asthma, there are two GWAS genes (IL13, PDE4D) that are explicitly targeted by asthma drugs, see table below. However, this enrichment is not significant p=0.7. We repeated this analysis across 35 MeSH diseases with known drug targets and 5 or more GWAS genes. Data is shown in Supplementary Table 1.

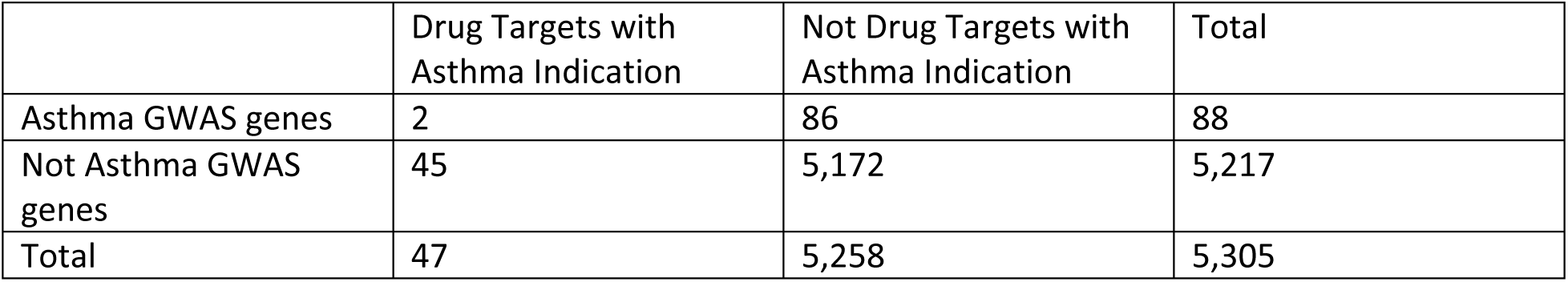

For each GWAS trait, we identified enriched pathways for “candidate effector genes” mapped to the GWAS loci. We corrected these pathway overlaps to ensure no more than one gene was used from any loci (Agarwal et al, 2006) (Supplementary Table 2). We then asked the question were the genes in these pathways enriched for drug targets for the disease matching the GWAS trait. Again, we built contingency tables as shown below for each disease.

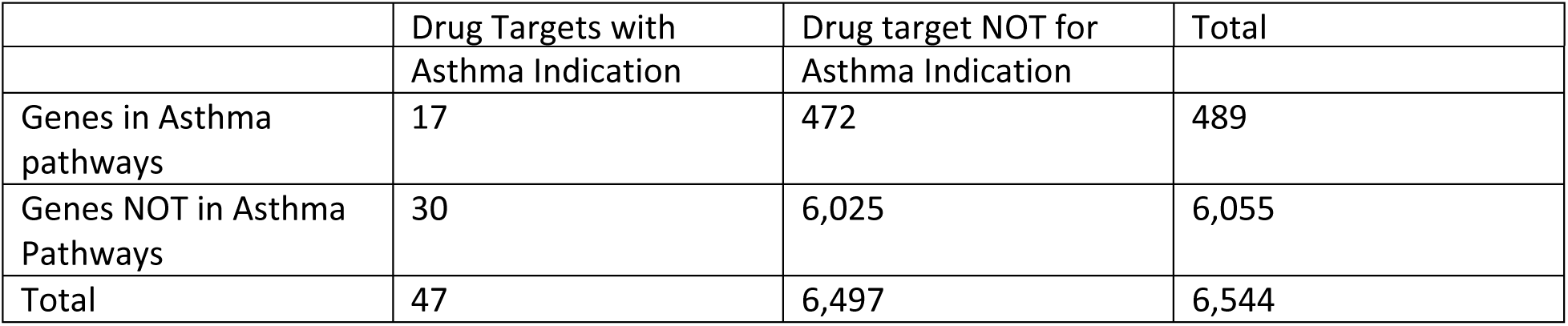

For example, Fisher’s exact test for asthma provides an enrichment p-value =1.8e-8 (p=2.18e-7 with Benjamini-Hochberg correction for FDR), suggesting that the pathways enriched for genes in asthma GWAS loci are also enriched for targets of drugs for asthma. The tractable genes from the 472 may in fact make good targets for asthma. These enrichments across 97 diseases are shown in Supplementary Table 3. Note that the number of diseases with 5 or more variants and known drug targets increased to 97 (from 35 in Supplementary Table 1) highlighting additional value of using all genes in each locus as potentially disease related.

## Acknowledgements

Authors would like to thank Andrew D. Rouillard, Daniel F. Simola, David N. Mayhew, and Philippe Sanseau for their valuable insights and discussions on this manuscript.

## Author Contributions

DJ, MMS, PA designed the study. DJ and PA performed the analysis. DJ, MMS, MH, PA interpreted the results and wrote the manuscript.

## Reference List

Agarwal P, Ghosh S, Hurle MR, Kabnick KS, Kumar VD, Liu L, Magid-Slav M, Mcallister PR, Reisdorf WC, Searls DB. (2006) Biological data set comparison method. Patent publication number US20070168135 A1.

Ailem M, Role F, Nadif M, Demenais F (2016) Unsupervised text mining for assessing and augmenting GWAS results. Journal of biomedical informatics 60: 252-259

Brodie A, Tovia-Brodie O, Ofran Y (2014) Large scale analysis of phenotype-pathway relationships based on GWAS results. PloS one 9: e100887

Cao C, Moult J (2014) GWAS and drug targets. BMC genomics 15 Suppl 4: S5

ClinicalTrials.gov. Efficacy and Safety of Ustekinumab, a Human Monoclonal Anti-IL-12/IL-23 Antibody, in Patients With Behçet Disease (STELABEC).

Cook D, Brown D, Alexander R, March R, Morgan P, Satterthwaite G, Pangalos MN (2014) Lessons learned from the fate of AstraZeneca’s drug pipeline: a five-dimensional framework. Nature reviews Drug discovery 13: 419-431

Hirschhorn JN (2009) Genomewide association studies‐‐illuminating biologic pathways. The New England journal of medicine 360: 1699-1701

Hurle MR, Nelson MR, Agarwal P, Cardon LR (2016) Trial watch: Impact of genetically supported target selection on R&D productivity. Nature reviews Drug discovery 15: 596-597

Lanczos C (1964) A precision approximation of the gamma function. Journal of the Society for Industrial and Applied Mathematics Series B Numerical Analysis 1: 86-96

Mooney MA, Nigg JT, McWeeney SK, Wilmot B (2014) Functional and genomic context in pathway analysis of GWAS data. Trends in genetics: TIG 30: 390-400

Nair JR, Moots RJ (2017) Behcet’s disease. Clinical medicine (London, England) 17: 71-77

Nelson MR, Tipney H, Painter JL, Shen J, Nicoletti P, Shen Y, Floratos A, Sham PC, Li MJ, Wang J, Cardon LR, Whittaker JC, Sanseau P (2015) The support of human genetic evidence for approved drug indications. Nature genetics 47: 856-860

Nikolsky Y, Kirillov E, Zuev R, Rakhmatulin E, Nikolskaya T (2009) Functional analysis of OMICs data and small molecule compounds in an integrated “knowledge-based” platform. Methods in molecular biology (Clifton, NJ) 563: 177-196

Pers TH, Karjalainen JM, Chan Y, Westra HJ, Wood AR, Yang J, Lui JC, Vedantam S, Gustafsson S, Esko T, Frayling T, Speliotes EK, Boehnke M, Raychaudhuri S, Fehrmann RS, Hirschhorn JN, Franke L (2015) Biological interpretation of genome-wide association studies using predicted gene functions. Nature communications 6: 58-90

Raychaudhuri S, Plenge RM, Rossin EJ, Ng AC, Purcell SM, Sklar P, Scolnick EM, Xavier RJ, Altshuler D, Daly MJ (2009) Identifying relationships among genomic disease regions: predicting genes at pathogenic SNP associations and rare deletions. PLoS genetics 5: e1000534

Sagar-Ouriaghli I, Lievesley K, Santosh PJ (2018) Propranolol for treating emotional, behavioural, autonomic dysregulation in children and adolescents with autism spectrum disorders. Journal of psychopharmacology (Oxford, England): 269881118756245

Sanseau P, Agarwal P, Barnes MR, Pastinen T, Richards JB, Cardon LR, Mooser V (2012) Use of genome-wide association studies for drug repositioning. Nature biotechnology 30: 317-320

Shen J, Song K, Slater A, Ferrero E, Nelson MR (2017) STOPGAP: a database for systematic target opportunity assessment by genetic association predictions. Bioinformatics (Oxford, England)

Spain SL, Barrett JC (2015) Strategies for fine-mapping complex traits. Human molecular genetics 24: R111-119

Sun J, Zhu K, Zheng W, Xu H (2015) A comparative study of disease genes and drug targets in the human protein interactome. BMC bioinformatics 16 Suppl 5: S1

Visscher PM, Wray NR, Zhang Q, Sklar P, McCarthy MI, Brown MA, Yang J (2017) 10 Years of GWAS Discovery: Biology, Function, and Translation. American journal of human genetics 101: 5-22

Wang K, Li M, Hakonarson H (2010) Analysing biological pathways in genome-wide association studies. Nature reviews Genetics 11: 843-854

